# *Klebsiella pneumoniae* OmpR facilitates lung infection through transcriptional regulation of key virulence factors

**DOI:** 10.1101/2023.07.31.550992

**Authors:** Axel B. Janssen, Vincent de Bakker, Rieza Aprianto, Vincent Trebosc, Christian Kemmer, Michel Pieren, Jan-Willem Veening

## Abstract

Bacteria must adapt to the stresses of specific environmental conditions to survive. This adaptation is often achieved by altering gene expression through two-component regulatory systems (TCSs). In Gram-negative bacteria, the response to environmental changes in osmolarity and pH are primarily mediated by the EnvZ/OmpR TCS. Although the functioning of EnvZ/OmpR has been well characterized in *Escherichia coli*, *Salmonella enterica*, and the *Yersinia* genus, the importance of EnvZ/OmpR TCS in the opportunistic human pathogen *Klebsiella pneumoniae* has been limitedly studied.

Here, we investigated the importance of EnvZ/OmpR in *K. pneumoniae* for fitness, gene regulation, virulence, and infection. Through the generation of a markerless *ompR*-deletion mutant, we show that overall fitness of *K. pneumoniae* is not impacted *in vitro*. Using dual RNA-seq of *K. pneumoniae* co-incubated with human lung epithelial cells we demonstrate that the *K. pneumoniae* OmpR regulon includes important virulence factors, but shows otherwise limited overlap with the regulons of other Gram-negative bacteria. In addition, we show that deletion of *ompR* in *K. pneumoniae* leads to a stronger antibacterial transcriptional response in human lung epithelial cells. Lastly, we show that OmpR is crucial for *K. pneumoniae* virulence and infection through a murine lung infection model.

As the adaptation of commensal bacteria to specific niches is mediated by TCSs, we show that EnvZ/OmpR plays a crucial role in successful lung infection, as well as in virulence. These results suggest that OmpR is an interesting target for anti-virulence drug discovery programs.

**Importance:** Bacteria use two-component regulatory systems (TCSs) to adapt to changes in their environment by changing their gene expression. In this study, we show that the EnvZ/OmpR TCS of *Klebsiella pneumoniae* plays an important role in successfully establishing lung infection, and virulence. In addition, we discern the transcriptional response that OmpR facilitates within this clinically relevant opportunistic pathogen, and within the host. This work suggests that *K. pneumoniae* OmpR might be a promising target for innovative anti-infectives.

## Introduction

*Klebsiella pneumoniae* is an important nosocomial pathogen due to the rapidly increasing rate of multidrug resistance. The spread of strains resistant to fluoroquinolones, third-generation cephalosporins, aminoglycosides, and increasing spread of resistance to last-line antibiotics such as carbapenemases and colistin, have limited treatment options (1, 2). This has made *K. pneumoniae* one of the pathogens with the highest burden of antimicrobial resistance deaths (3). Due to this multidrug resistance, and lack of development of novel antibiotics *K. pneumoniae* has been named by the World Health Organization as a critical priority pathogen for the development of novel antibiotics (4).

*K. pneumoniae* can asymptomatically colonize the upper respiratory tract, skin, and digestive tract in healthy individuals. However, the asymptomatic colonization of these niches by *K. pneumoniae* may subsequently lead to infections such as lung infections, soft tissue infections, wound infections, and urinary tract infections (5). The populations that are particularly at risk of *K. pneumoniae* infections are infants, the elderly, the immunocompromised, and hospitalized patients (5, 6). Persistent asymptomatic niche colonization by potential pathogens is mediated by adaptation to niche-specific conditions. Two-component regulatory systems (TCSs) mediate the adaptation of bacteria in response to changing environmental factors through adaptation of gene expression. TCSs are composed of a membrane-bound sensor histidine kinase, and a cytosolic transcriptional response regulator (7, 8). The membrane-bound sensor kinase will activate through *trans*-autophosphorylation in response to a specific environmental signal (8, 9). Next, the phosphorylated sensor histidine kinase will transfer its phosphoryl-group to the response regulator, leading to conformational changes within the response regulator. The phosphorylated response regulator will then bind to specific promoter sequences, resulting in changed gene expression and an increased survival under the specific environmental conditions (10–12). TCSs thus play an important role in persistent colonization, and subsequent infection, by potential pathogens like *K. pneumoniae*, in diverse niches.

The EnvZ/OmpR TCS is involved in the adaptive response to environmental changes in both osmolarity and pH (13–15). EnvZ is the sensor histidine kinase part of this TCS. In the presence of specific activation signals, the EnvZ dimer will phosphorylate the OmpR transcriptional response regulator. Two phosphorylated OmpR molecules will bind to the consensus target sequences, resulting in an altered gene expression (11, 16). The role of EnvZ/OmpR in gene regulation, virulence, colonization, and infection, has been well studied in several bacterial species, like *Escherichia coli* (including uropathogenic, and adherent invasive strains) (13, 17–22), *Yersinia* spp. (23–25), *Salmonella enterica* (26, 27), *Shigella flexneri* (28), and *Acinetobacter baumannii* (29), often suggesting that EnvZ/OmpR plays a key role in pathogenesis. The number of studies in *K. pneumoniae* have however, been limited.

Recently, it has been shown that *K. pneumoniae* ATCC 43816 deficient in OmpR has reduced virulence in a mouse lung-infection model (30). This reduction in virulence has been suggested to be the result of the loss of hypermucoviscovity because of the loss of OmpR. Although OmpR seems to be involved in regulation of hypermucoviscocity *in vitro*, the transcriptional changes upon deletion of *ompR* in a host-pathogen setting have not been elucidated. Here, we studied the functioning of the EnvZ/OmpR TCS in *K. pneumoniae* ATCC 43816 on fitness, gene expression, and infection and virulence. We utilized dual RNA-seq (31) to elucidate the role of OmpR as a transcriptional regulator in early host-pathogen interaction between *K. pneumoniae* and human lung epithelial cells. In addition, we confirm the role of OmpR in infection and virulence through a lung infection mouse model.

## Materials and Methods

### Ethical statement

Animal experiments were performed by Aptuit (today Evotec; Verona, Italy). All experiments involving animals were carried out in accordance with directive 2010/63/EU of the European Union governing the welfare and protection of animals, implemented by Italian Legislative Decree number 26/2014, and in accordance with Aptuit’s policy on the care and use of laboratories animals. All animal studies were reviewed by an Animal Welfare Body (Aptuit) and approved by Italian Ministry of Health.

### Bacterial strains, growth conditions, and mutant construction

*K. pneumoniae* ATCC 43816 WT was obtained from the American Type Culture Collection (ATCC; Manassas, Virginia, United States of America). *K. pneumoniae* ATCC 43816 WT and derivate strains were cultured in lysogeny broth (LB) without antibiotics, at 37°C with agitation, unless otherwise noted. The deletion of *ompR* was performed through an allelic exchange method using a combination of positive selection (selection through sodium tellurite), and negative selection (selection through SceI restriction endonuclease), as described before (18, 32).

### Determination of maximum specific growth rate

Maximum specific growth rates were determined by measuring growth in a Tecan Infinite M Plex plate reader (Tecan, Männedorf, Switzerland). Overnight cultures in LB, or RPMI1640 without phenol red (Gibco, Life Technologies, Bleiswijk, the Netherlands) supplemented with 1% fetal bovine serum (FBS; Gibco), were used to inoculate 200 μl of the same medium 1:1000. Samples were incubated at 37°C with agitation. Growth was observed by measuring the absorbance at 595 nm (Abs_595_) every 10 minutes. Maximum specific growth speed was determined by calculating the maximum increase in natural logarithm transformed absorbance values between two adjacent timepoints, divided by time between timepoints in hours: 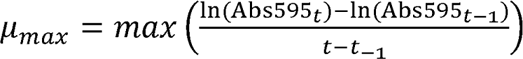

### Determination of minimal inhibitory concentration

Minimal inhibitory concentrations (MICs) were determined by microbroth dilution method in cation-adjusted Mueller-Hinton broth according to Clinical and Laboratory Standards Institute (CLSI) guidelines, except for the MIC of ertapenem from the ATCC 43816 WT strain, that was determined through a Vitek 2, using an AST-GN69 card (BioMérieux, Marcy-l’Étoile, France). The CLSI quality control strain *E. coli* ATCC 25922 was included in the microbroth dilution MICs as control.

### *In vitro* infection studies

The human type II lung epithelial cell line A549 (ATCC CCL-185) was cultured in DMEM/F12 with GlutaMAX (Life Technologies) supplemented with 10% FBS (FBS; VWR, Amsterdam, The Netherlands) under controlled conditions (37°C, 5% (v/v) CO_2_). Prior to the start of the infection experiment, epithelial monolayers were incubated for 10 days after confluence in medium without antibiotics. Confluent monolayers of A549 were co-incubated with either *K. pneumoniae* WT or Δ*ompR* strain at a multiplicity of infection of 10. The infection experiment was performed in RPMI1640 medium without phenol red with 1% FBS. A spin infection was performed (2000□×□*g*, 5 min, 4°C) to promote the contact between bacterial and epithelial cells immediately after the addition of the bacterial suspension. The samples were co-incubated at 37°C and 5% CO_2_ for 2 hours to mimic early infection, after which the total RNA from both the *K. pneumoniae* and A549 cells was isolated.

### RNA isolation

Total RNA was simultaneously isolated from the harvested mixture of co-incubated *K. pneumoniae* and A549 cells. To minimize transcriptional changes during sample handling, we did not separate the cellular mixture of bacterial and human cells before isolation. To prevent protein-dependent RNA degradation, we treated the cellular mixture with a saturated buffer solution of ammonium sulfate (pH 5.2, 700 g/L (NH_4_)_2_SO_4_), also containing 20 mM EDTA and 25 mM sodium citrate (33). Three parts of saturated solution of ammonium sulfate were mixed to one part of infection medium. The suspension was vigorously pipetted to ensure the complete mixing of the saturated buffer solution and infection medium. After scraping of the adherent host cells the suspension was incubated further at room temperature for 5 minutes. The suspension was collected and centrifuged at full speed (20 min, 4°C, 10,000□×□*g*). The supernatant was removed, and the cell pellet was snap frozen in liquid nitrogen.

A PCR tube full of sterile, RNase-free 100 μm glass beads (BioSpec, Bartlesville, Oklohoma, United States of America) was mixed with 50 μl 10% SDS and 500 μl phenol-chloroform in a 1.5 ml screw cap tube. The frozen cell pellet was resuspended in TE solution (10 mM Tris-HCl, 1 mM Na_2_DTA, pH 8.0, in DEPC treated milliQ H_2_O). The suspension was added into the screw cap tube and bead-beaten in three cycles of 45 seconds each. Tubes were immediately placed on ice and centrifuged at full speed at 4°C to separate organic and aqueous phases. After removing the aqueous phase, back-extraction was performed on the organic phase for optimal RNA yield. Phenol was further depleted from the aqueous phase by another round of chloroform extraction. After vigorous vortexing, the mixture was again centrifuged at full speed, 4°C. The aqueous phase was transferred into a fresh eppendorf tube and nucleic acids were precipitated by addition of 50 µl 3M NaOAc and 1 ml of cold isopropanol, and vigorous mixing. The mixture was incubated for at least 30 minutes at -20°C before pelleting by centrifugation (full speed, 4°C). Supernatant was carefully removed, and the nucleic acid pellet washed twice by resuspension in ice-cold 75% ethanol before re-pelleting (full speed, 4°C). The pellet was air-dried before DNase treatment.

DNase treatment was performed using recombinant RNase-free DNase I (Roche, Basel, Switzerland) according to the manufacturer’s protocols for 1 hour, at room temperature. To remove DNase and gDNA-derived nucleotides, phenol-chloroform extraction, chloroform extraction, isopropanol precipitation, and ethanol washing were performed as previously mentioned. Total RNA was resuspended in 30 µl TE buffer. The quantity and quality of total RNA was estimated by Nanodrop and a 1% bleach gel was employed to assess the presence of genomic DNA and rRNA bands (23S; 2.9 kbp, and 16S; 1.5 kbp) (34).

### Library preparation and sequencing for dual RNA-seq

The quality of the total RNA isolated was checked using chip-based capillary electrophoresis. Samples were simultaneously depleted from human and bacterial ribosomal RNAs by dual rRNA-depletion as previously described (33). Reverse stranded cDNA library preparation was performed with the TruSeq® Stranded Total RNA Sample Preparation Kit (Illumina, San Diego, California, United States of America) according to the manufacturer’s protocol. Single-ended sequencing was performed on single lane of a High Output Flowcell in an Illumina NextSeq 500 (Illumina) instrument.

### RNA-Seq data analysis

Quality of raw sequencing reads was checked using FastQC (v0.11.9; available at https://www.bioinformatics.babraham.ac.uk/projects/fastqc/; Babraham Bioinformatics, UK). Reads were trimmed for adapter sequences from the TruSeq3-SE library preparation kit, minimal quality of starting and trailing nucleotides (Phred score of 20), a cutoff of an average Phred score of 20 in a 5-nucleotide sliding window, and length (minimum length 50 nucleotides) using Trimmomatic (v0.39) (35). The quality of trimmed reads was confirmed using FastQC.

Quality filtered reads were aligned to a chimeric sequence created by concatenating the circular genome of *K. pneumoniae* ATCC 43816 (Genbank accession GCA_016071735.1) (36), and the human genome (Genbank accession GCA_009914755.4) (37). The corresponding annotation files were downloaded from the same services. First, indexing of the chimeric genome was performed using RNA-STAR with the following options: --alignIntronMax set to 1, and --sjdbOverhang set to 84. Then, alignment was performed by using RNA-STAR (v2.7.10a) (38) with default settings. The aligned reads were summarized through featureCounts (v2.0.1) (39) according to the chimeric annotation file in reverse stranded (-s 2), multimapping (-M), fractionized (--fraction) and overlapping (-O) modes. The single-pass alignment onto the chimeric genome was selected to minimize the false discovery rate. However, due to this approach, we had to adjust the summarizing process, considering the overlapping nature of bacterial genes and their organization into operon structures. In case of conflicting gene annotations between species, the naming in these files were deemed leading.

The counts of the aligned reads of the host and pathogen libraries were analyzed separately in R (v4.1.1). Differential expression analysis was performed with DESeq2 v1.34.0 (40). For principal component analyses (PCA), counts were normalized with a blind rlog transformation. Since coverage for the bacterial data set was high and well-saturated, we used stricter parameters than the default for hypothesis testing: 1.5× depletion or enrichment of mRNAs (|log_2_FC|>log_2_(1.5)) with P_adj_<0.05 as significance threshold. As the host data set was less well-saturated, we left the log_2_FC parameter at its default (|log_2_FC|>0: any difference) to look for more subtle effects, but restricted the significance threshold to P_adj_<0.01 to retain high confidence in called hits. GO term enrichment and gene set enrichment analyses were performed with clusterProfiler (v4.2.2) (41) using the log_2_FC values from DESeq2, shrunk with the apeglm (42) method as implemented in that package. GO terms were clustered with the aPEAR package (43).

### *In vivo* lung infection studies

Male CD-1 mice (15 animals per bacterial strain and inoculum) were infected intranasally with 10^5^ or 10^6^ colony forming units (CFU) (50 µl) of exponential phase cultured *K. pneumoniae* ATCC 43816 WT or Δ*ompR*. Five animals per group were euthanized using CO_2_ at 2-, 24-, and 48-hours post-infection. Aseptically sampled lungs and blood were assessed for bacterial titers through CFU counting. Survival rate, clinical severity scores and body weight were recorded during the experimental phase. The score system from 0 (normal) to 5 (death) was built by Aptuit on the basis of internal experience and knowledge of the specific model about development of the infection.

### Statistical analyses

Except for host differential expression analysis, statistical significance was defined as a p-value of <0.05 for all tests. Significance was defined as a p-value ≤ 0.05 (*), p ≤ 0.01 (**), p ≤ 0.001 (***), or p ≤ 0.0001 (****). Statistical analyses other than RNA-seq analyses were performed using GraphPad Prism 6 software (GraphPad Software, San Diego, CA, USA) and R (v4.0.3).

## Results

### Deletion of *ompR* does not impact fitness of *K. pneumoniae* ATCC 43816 *in vitro*

As a limited number of studies on the role of EnvZ/OmpR in *K. pneumoniae* have previously been performed, we first aimed to characterize the effects of the deletion of *ompR* in *K. pneumoniae* ATCC 43816 *in vitro*. We constructed a markerless deletion mutant through an allelic exchange method using a combination of positive and negative selection (32). To assess the effects of the deletion of *ompR* on fitness, we determined the growth kinetics of *K. pneumoniae* ATCC 43816 and the Δ*ompR* strain. We observed that the deletion of *ompR* did not lead to an impairment of overall growth kinetics (Figure 1A). To quantify the growth kinetics in more detail, we determined the maximum specific growth rate, which is an accepted proxy for overall fitness of a bacterial strain (44). Through tracking of the growth, we observed that the maximum specific growth rate of the Δ*ompR* strain was not affected compared to the WT strain, in either LB, or RPMI1640 supplemented with 1% FBS (Figure 1B). Thus, a strain deficient in OmpR does not seem to have any growth defects *in vitro*.

**Figure 1.**
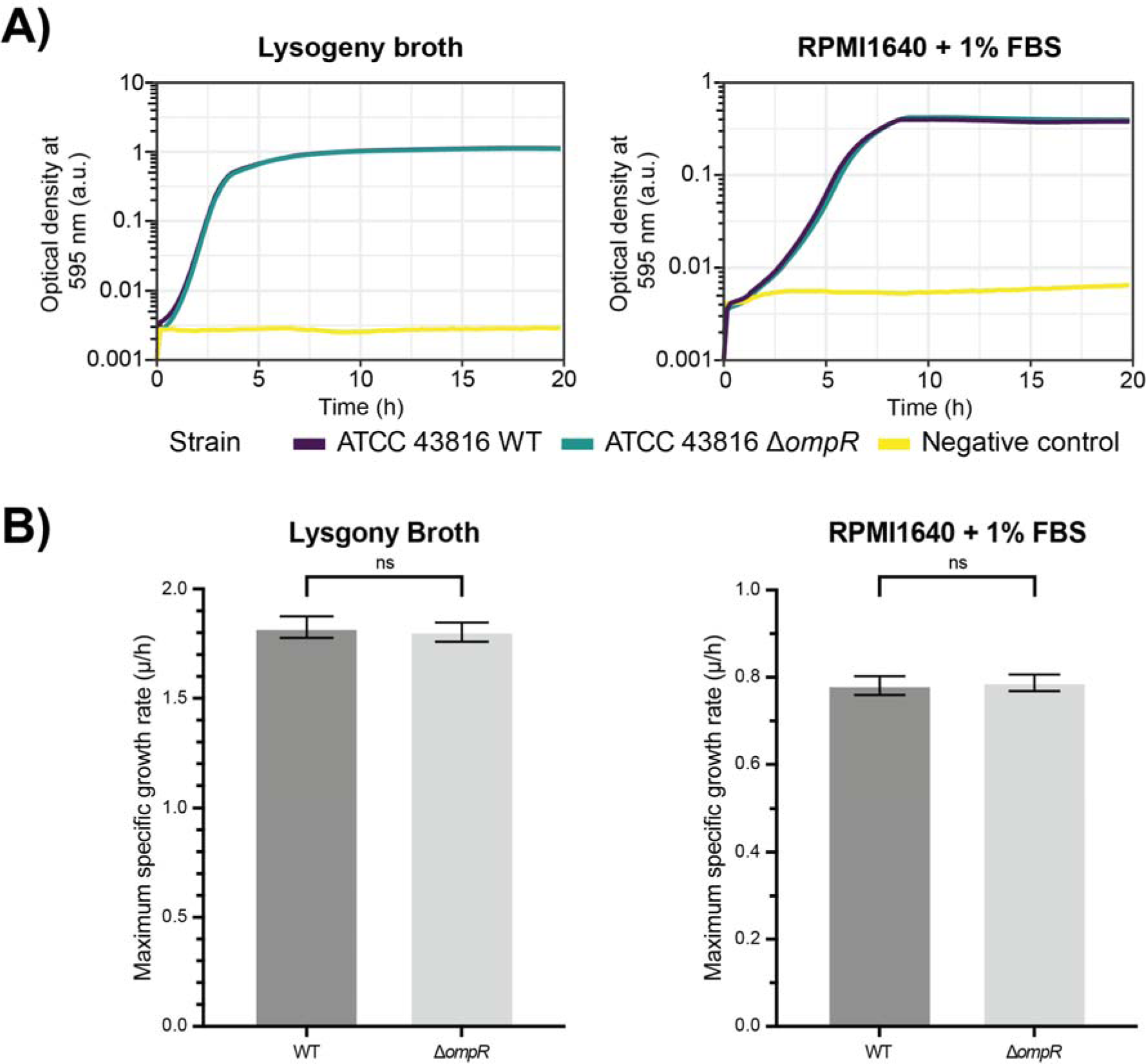
Deletion of *ompR* in *K. pneumoniae* ATCC 43816 does not lead to a major fitness defect *in vitro*. **A)** Determination of the growth kinetics of ATCC 43816 WT and ATCC 43816 Δ*ompR* showed that the deletion of *ompR* does not lead to changed growth kinetics in either LB, or RPMI1640 complemented with 1% FBS. **B)** Determination of the maximum growth speed as a measure of fitness showed that deletion of *ompR* does not lead to a reduced fitness in either LB, or RPMI1640 complemented with 1% FBS. Statistical significance was determined using an unpaired T-test.

In addition, we assessed whether the deletion of *ompR* in *K. pneumoniae* ATCC 43816 leads to changes in antibiotic susceptibility. We subjected the WT and the Δ*ompR* strain to standardized minimal inhibitory concentration (MIC) determinations of several types of antibiotics. We observed that the susceptibility of the Δ*ompR* strain did not change more than 2-fold, compared to the WT strain, for all but the tetracycline molecules. The MIC values were 4-fold higher for both tetracycline (1 µg/ml to 4 µg/ml), and minocycline (0.5 µg/ml to 2 µg/ml) for the Δ*ompR* strain compared to the WT strain (Supplemental Table S1). These increases did not however make the Δ*ompR* strain phenotypically resistant to the tested tetracyclines, according to the breakpoints provided by the CLSI.

### Limited similarity in OmpR regulon between *K. pneumoniae* and other species during early host-pathogen interaction

Because OmpR is the transcriptional regulator part of the EnvZ/OmpR TCS, we questioned what consequences the *ompR* deletion has on gene expression of *K. pneumoniae* ATCC 43816. To investigate the gene expression, we utilized dual RNA-seq. The simultaneous genome-wide profiling of host and pathogen transcriptional responses using deep sequencing (i.e. dual RNA-seq) offers limited technical bias, and higher efficiency compared to conventional approaches (e.g. single species approach, or array-based methods) (31).

As *K. pneumoniae* is a well-known lung pathogen, the pathogen was co-incubated with a monolayer of human A549 lung epithelial cells, which have been widely used as a model for respiratory infections (45, 46). The lung epithelial cells thus resemble a relevant substrate for the pathogen for the study of early host-pathogen interactions. We cultured the A549 lung epithelial cells without antibiotics to exclude any effects due to the residual presence of antibiotics, or their downstream effects; for example, the presence of bacterial fragments (e.g. intracellular) and their effects on the host response. As the main interest in this experiment was to assess the early host-pathogen interaction, we selected the timepoint of two hours after the start of co-incubation (2 hpi) of the A549 and the *K. pneumoniae* cells as the endpoint of this infection model.

For the dual RNA-seq analysis, three samples of A549 lung epithelial cells infected with ATCC 43816 WT, and two samples of A549 lung epithelial cells infected with the ATCC 43816 Δ*ompR* mutant strain were available (one Δ*ompR* replicate was unfortunately lost during sample preparation). Global analysis of the dual RNA-seq libraries of early infection showed that an average of 54.7 million (range 36.2 million - 74.0 million; Supplemental Figure S1A) reads mapped to the hybrid host-pathogen genome. Most aligned reads mapped onto the human genome (average 65.3%, range: 61.3 - 68.9%; Supplemental Figure S1B). However, higher saturation and read coverage was observed for the bacterial genes (Supplemental Figure S1A). Normalized read counts of the RNA-seq data set showed that replicate infections with either WT or Δ*ompR* were reproducible (Pearson’s r >0.995; Supplemental Figure S1C) and that most of the transcriptional variation for either organism was attributable to the deletion of *ompR* (Supplemental Figure 1D).

Within the pathogen data set, we observed that deletion of *ompR* in the genome of *K. pneumoniae* ATCC 43816 led to a statistically significantly 1.5× lower (P_adj_<0.05) expression of 12 genes, whilst 11 genes were statistically significantly 1.5× higher (P_adj_<0.05) expressed (Figure 2, Table 1, Supplemental Table S2). Several of these genes have been previously reported to be members of the OmpR regulon in other bacterial species. Genes encoding outer membrane porins *ompK35* and *ompC* (21), but also *dtpA* (20), encoding a di-/tripeptide permease, were lower expressed. We also observed higher expression of *mglB* (27), a galactose/glucose transporter. These genes can play a role in maintaining proper intracellular osmolarity in the bacterial cells by regulating transport of their specific substrates. We also observed a lower expression of *mrkA* (47), encoding a fimbrial type 3 subunit.

**Figure 2.**
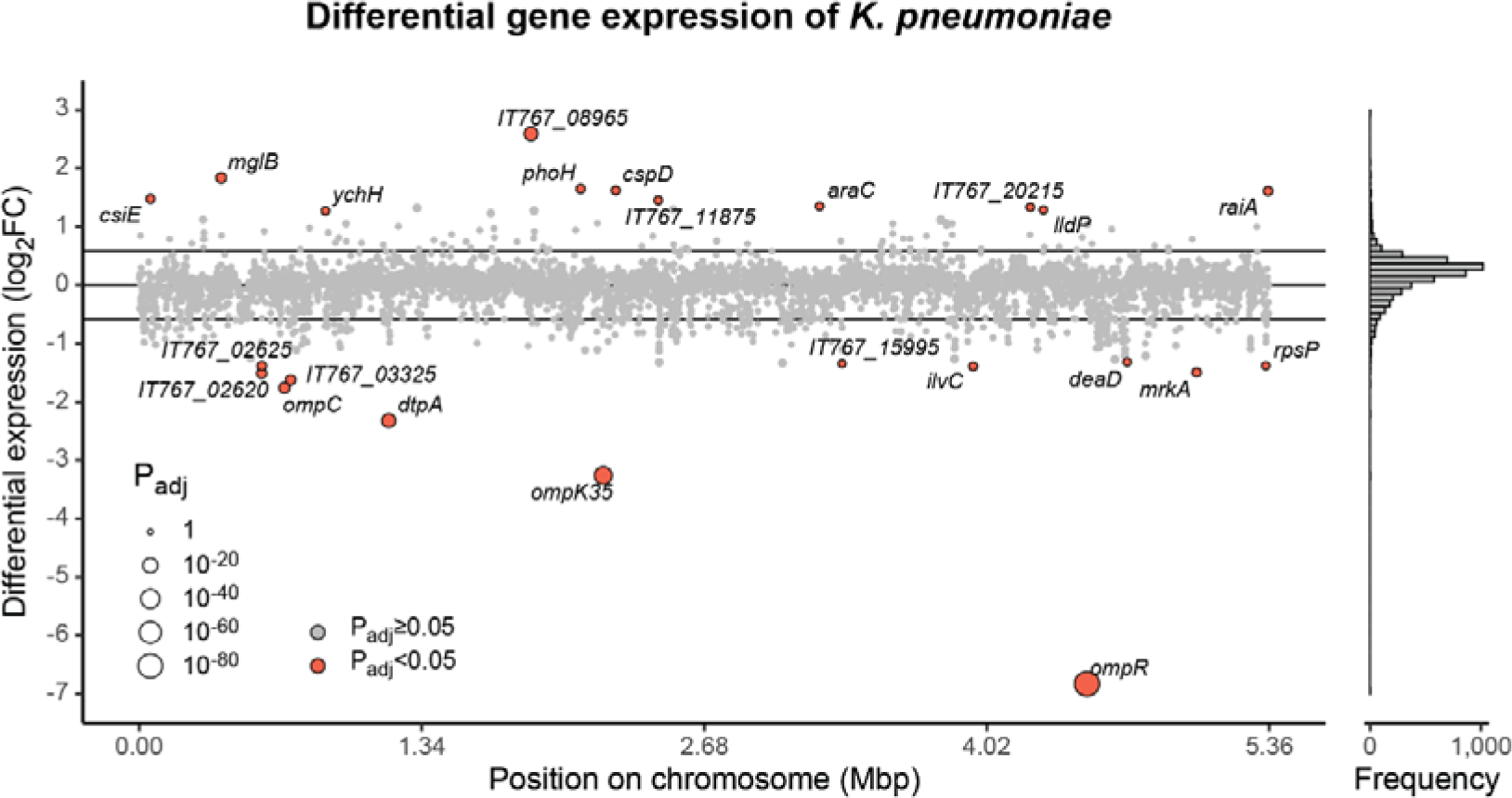
OmpR-mediated differential expression in *K. pneumoniae* co-incubated with human epithelial cells. Fold-changes (FC) in expression of *K. pneumoniae* ATCC 43816 genes in the Δ*ompR* strain relative to WT on a log_2_ scale. Point size indicates adjusted p-values on an inverse log_10_ scale and color indicates significance, compared to a 1.5-fold change in expression (horizontal lines). Gene names of significantly differentially expressed genes are indicated if annotated, locus tags are given otherwise. The histogram indicates the distribution of log_2_FC values to illustrate point density.

**Table 1.**
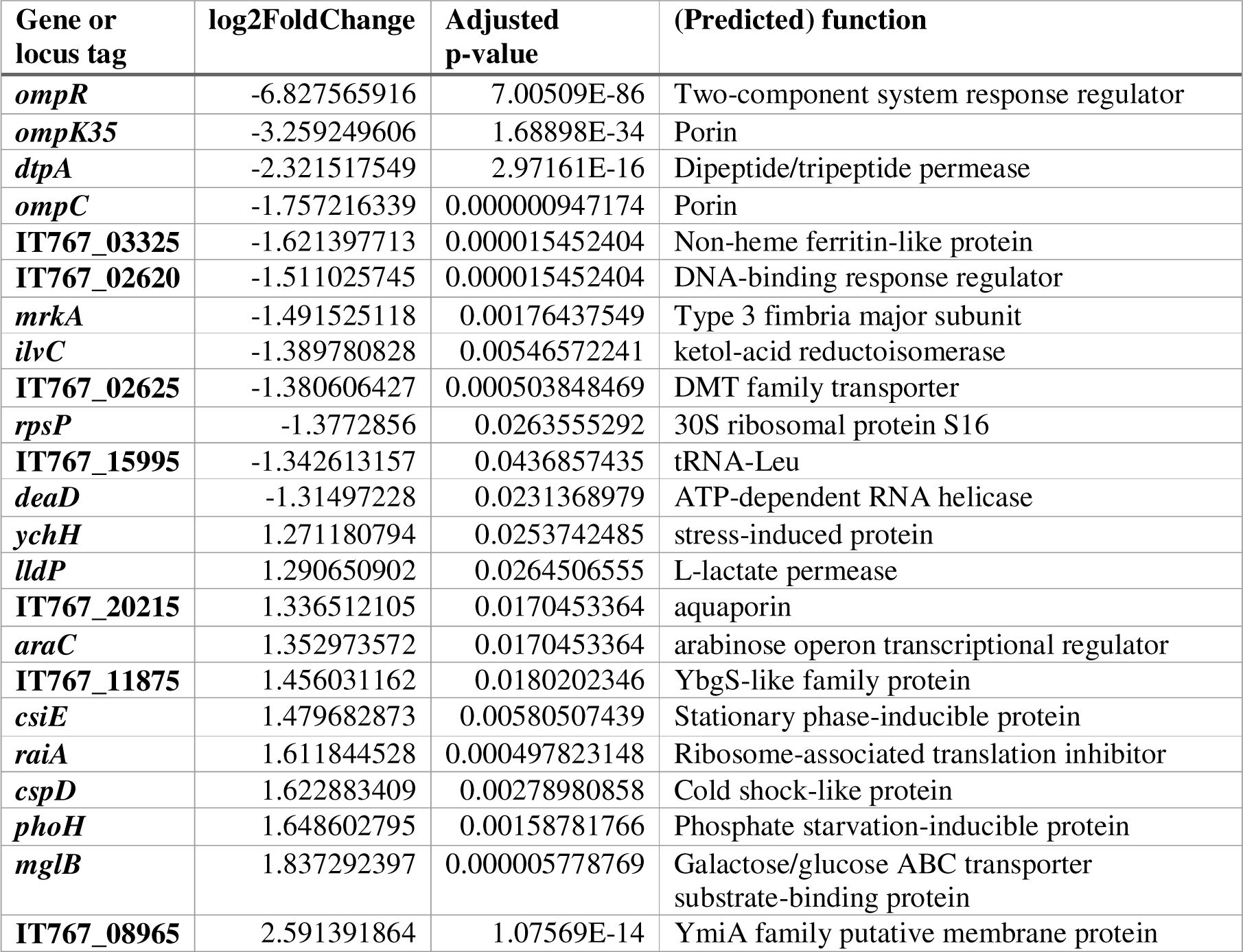
Genes significantly differentially expressed in *K. pneumoniae* Δ*ompR* compared to WT, when co-incubated with a monolayer of human A549 lung epithelial cells. Genes that are significantly differentially expressed (having an absolute log_2_FoldChange (log_2_FC) >log_2_(1.5), and an adjusted p-value < 0.05) in the Δ*ompR* strain compared to the WT strain. For each gene, the function, the adjusted p-value, and log_2_FoldChange are given. For genes that have not been named, the predicted function (as per the published annotation) is given.

Outside of the genes previously described to be in the OmpR regulon, we observed several other genes to be statistically significantly differentially expressed. Amongst others, IT767_03325, encoding a non-heme ferritin-like protein, and homologue of FtnB of *Salmonella* species, was lower expressed. In addition, we observed that IT767_02620, (previously renamed to *rmpC*), involved in regulation of the biosynthesis of capsular polysaccharides (48), was lower expressed as well. Higher expression was observed for IT767_20215, predicted to encode a porin involved in glycerol uptake, and *lldP*, encoding a L-lactate permease. These last two could impact the maintenance of proper intracellular osmolarity.

In an attempt to identify the preferential DNA sequence bound by OmpR, we compared the sequences (500 bp) immediately preceding the genes that were differentially expressed in our setup. From this analysis, no conclusive consensus DNA sequence could be identified (data not shown).

### *K. pneumoniae* OmpR influences anti-bacterial responses by human lung epithelial cells

In addition to the pathogen response to deletion of *ompR* within the *in vitro* infection model, we analyzed the transcriptional response on the host side to study OmpR-specific host responses during interaction with *K. pneumoniae*. In the human data set, out of 57816 annotated genes in the assembly of the human genome, 2786 genes were filtered out because no reads had mapped to the feature (Supplemental Table S3). A further 19153 genes were filtered out due to a low normalized read count (Supplemental Figure S1A), leaving 35877 genes (62%) for which differential expression was assessed (40).

Of these 35877 genes, we observed that 44 genes were significantly lower expressed (|log_2_FC|<0, P_adj_<0.01) in the cells exposed to the ATCC 43816 Δ*ompR* bacteria, compared to the cells exposed to the WT bacteria. A large portion of these lower expressed genes encode non-coding RNAs; mostly small nucleolar RNAs (snoRNAs), but also small nuclear RNAs (RNU) (Figure 3A, Supplemental Table S3). In addition to the non-coding RNAs, among the genes most negatively affected in expression in the mutant-exposed cells were multiple genes encoding ribosomal proteins (e.g. *RPSAP8*, and *RPS27P13*) (Figure 3A). In contrast, we found that no genes were significantly higher expressed (|log_2_FC|>0, P_adj_<0.01) in the lung epithelial cells exposed to ATCC 43816 Δ*ompR* bacteria compared to the lung epithelial cells exposed to the WT bacteria (Figure 3A, Supplemental Table S3).

**Figure 3.**
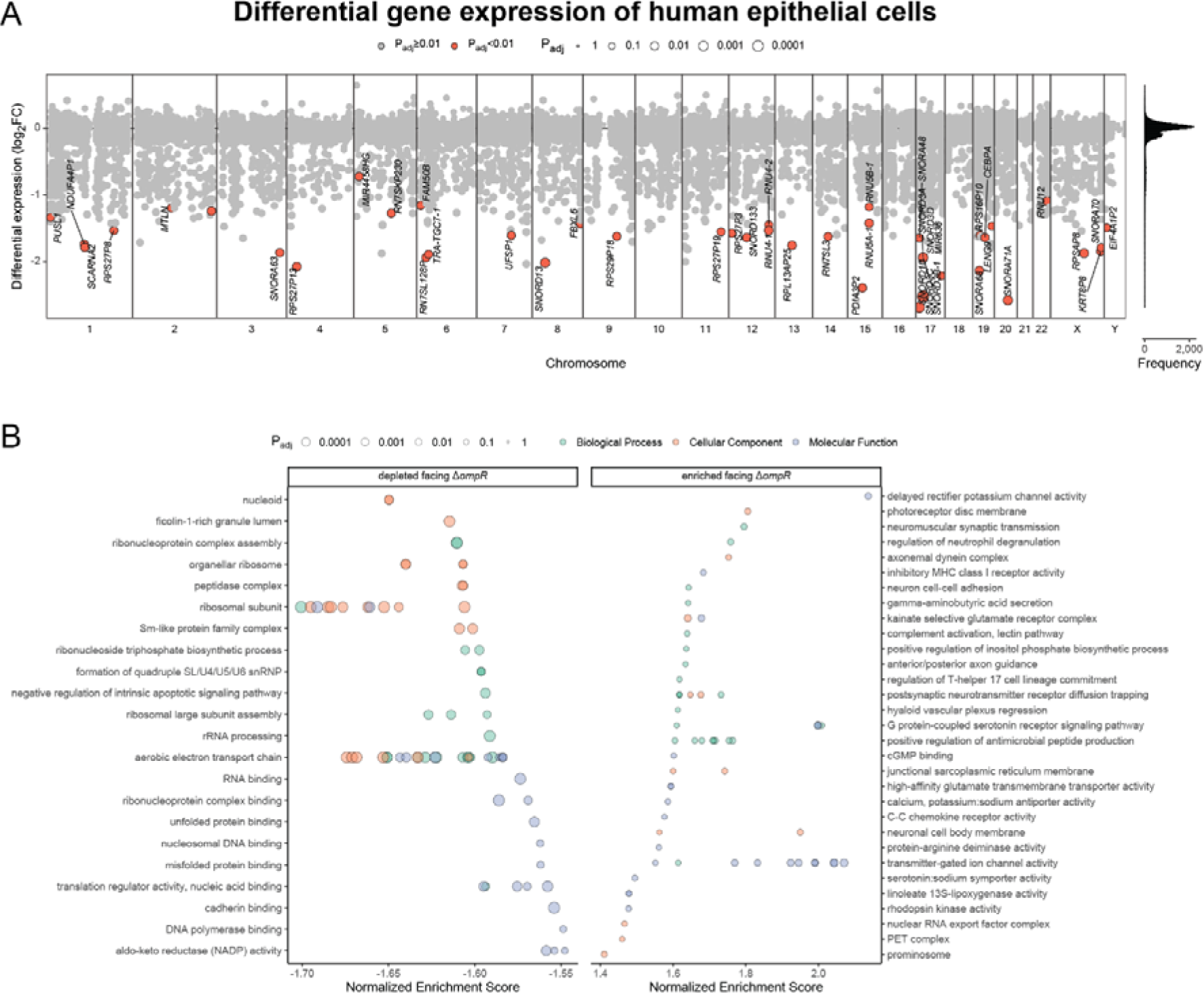
Transcriptional response of human A549 lung epithelial cells in response to co-incubation with *K. pneumoniae* ATCC 43816 WT or Δ*ompR* cells. **A)** Differential expression over all loci per chromosome (1-22, X, Y) of cells challenged with the Δ*ompR* strain relative to the WT strain. Significantly differentially expressed genes (|log_2_FC|>0, P_adj_<0.01) depicted in orange and with corresponding gene names if annotated. The histogram shows the distribution of the log_2_FC values to indicate point density. **B)** GO term gene set enrichment analysis (GSEA) of all log_2_FC values. GO terms were clustered with the aPEAR package. Shown are the top 25 GO terms with the highest and lowest normalized enrichment scores with P_adj_<0.1 for each data base (Biological Process, Cellular Component and Molecular Function).

To better understand the patterns in the differentially expressed genes between the cells exposed to ATCC 43816 WT cells, and those exposed to ATCC 43816 Δ*ompR* cells, we used a Gene Ontology (GO) term enrichment analysis. By using a GO term enrichment analysis, the RNA-seq data can be assessed for an entire ontology (either biological processes, molecular functions, and/or cellular components), instead of on an individual gene level. When analyzing only those genes that were significantly differentially expressed, the GO term enrichment analysis confirmed that RNA processing pathways were significantly downregulated in lung epithelial cells exposed to ATCC 43816 Δ*ompR* cells (Supplemental Table S4, Supplemental Figure 2).

However, as subtle (non-significant) changes in the expression of the genes that together constitute a biological process may sum up to significantly influence a process together, we also performed GO term gene set enrichment analysis (GSEA) using the log_2_FC values of all 55030 genes for which a fold change could be calculated. In addition to the significantly lower expression of RNA processing genes as identified before, this analysis also identified relative downregulation of gene sets related to household processes such as ribosomal functioning and respiration in the mutant-exposed cells (Figure 3B, Supplemental Table S5). In contrast to the GO terms describing ribosomal functioning, RNA processing, and respiration, GO terms involved in antimicrobial peptide production, innate immune responses, and neuronal communication were more highly expressed in the cells challenged with mutant bacteria (Figure 3B, Supplemental Table S5).

Through dual RNA-seq in an *in vitro* host-pathogen model, we identified that deletion of *ompR* in *K. pneumoniae* ATCC 43816 leads to lower expression of several genes involved in biogenesis of membrane-related structures such as *ompC* (annotated as *ompS1* in *Salmonella* species) and *ompK35*, but also of *mrkA* and IT767_02620 (*rmpC*). These genes, or the products that they regulate, have been previously described to be associated with virulence in diverse bacterial species (5, 19, 47, 49). In addition, we observed that the lung epithelial cells that were exposed to the mutant strain had higher expression of pathways involved in antimicrobial responses. These results, together with previous reports that OmpR is an important virulence factor in closely related bacterial species (18, 24, 29), suggest that the deletion of *ompR* in *K. pneumoniae* could also lead to a reduction in virulence.

### Deletion of *ompR* reduces virulence and infection of *K. pneumoniae* ATCC 43816

To assess the effects of the deletion of *ompR* on the virulence of *K. pneumoniae* ATCC 43816, and its ability to establish an infection in the lungs, male CD-1 mice were intranasally infected with exponential phase WT or Δ*ompR* bacteria.

The intranasal inoculation with 1.92 × 10^6^ CFU of the WT strain resulted in mean bacterial lung titers of 1.59 × 10^6^ and 9.75 × 10^6^ CFU at 2- and 24-hours post-infection (hpi) respectively (Figure 4A), and in 100% mortality at 48 hpi (Figure 4B). Inoculation with 1/10^th^ of that dose resulted in mean bacterial titers of 8.97 × 10^4^, 3.51 × 10^7^, and 1.08 × 10^8^ CFU per lung at 2-, 24-, and 48 hpi. In addition, decreased clinical severity scores and mortality at 48 hpi were observed compared to the higher inoculum (Figure 4B). In contrast to inoculation with WT cells, intranasal inoculation with 10^6^ CFU Δ*ompR* strain resulted in mean bacterial lung titers of 5.80 × 10^6^, 3.67 × 10^4^, and 6.63 × 10^2^ CFU at 2, 24, and 48 hpi respectively (Figure 4A). Inoculation with the lower dose of the Δ*ompR* strain resulted in mean bacterial lung titers of 2.22 × 10^5^, 4.00 × 10^3^, and 1.06 × 10^4^ CFU at 2, 24, and 48 hpi. None of the animals infected with the Δ*ompR* strain died. Mice infected with the Δ*ompR* mutant strain had decreased clinical severity scores during the 48 hours of monitoring post-infection, compared to the mice infected with the WT strain (Figure 4B).

**Figure 4.**
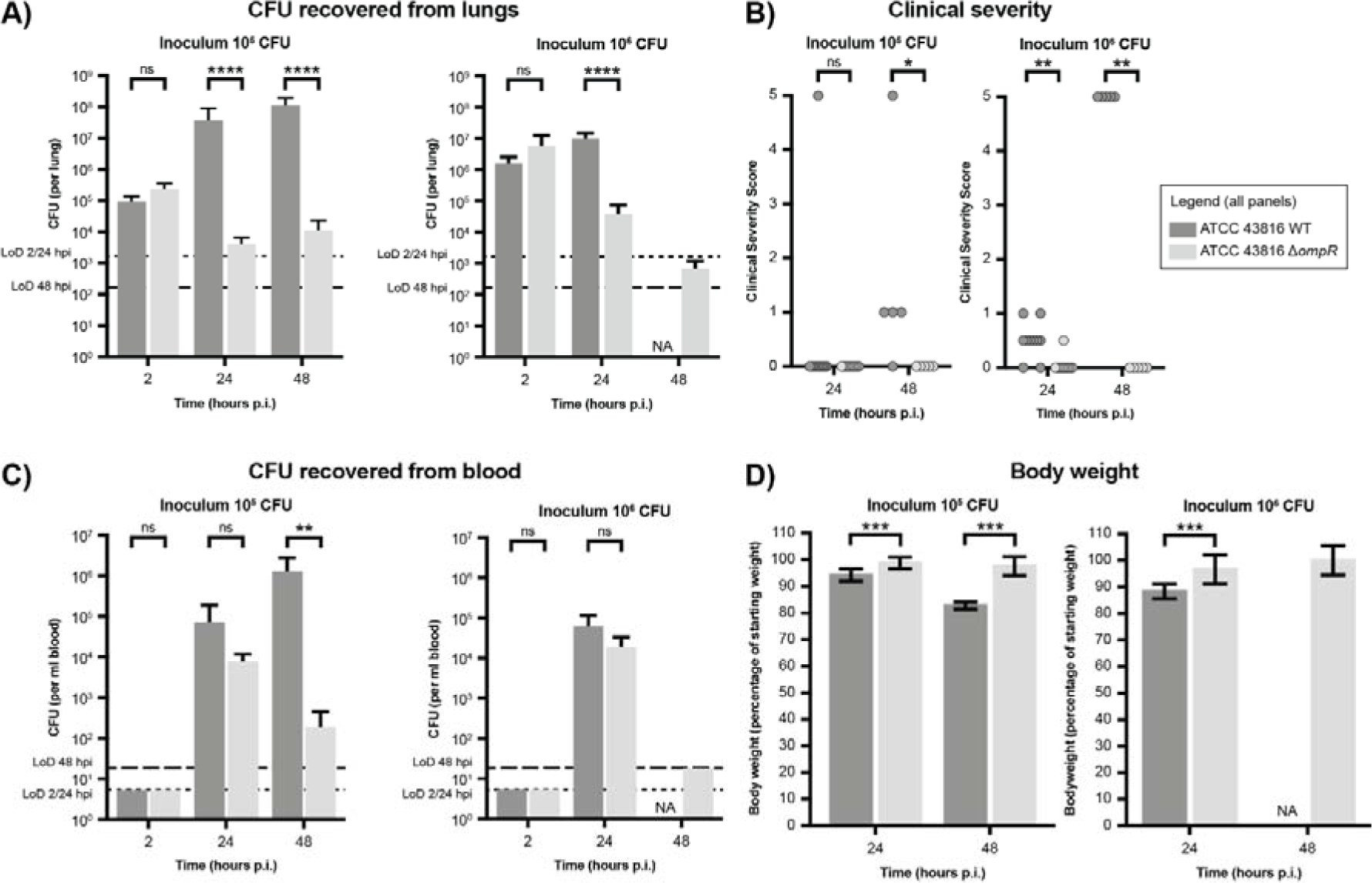
Deletion of *ompR* in *K. pneumoniae* leads to decreased bacterial loads in a *K. pneumoniae* lung infection model. CD-1 mice (15 per group) were infected intranasally with *K. pneumoniae* ATCC 43816 WT or the isogenic Δ*ompR* mutant strain. **A)** The lungs were collected for bacterial titer determination. For the group infected with high inoculum of WT cells, no data was gathered at 48 hpi, because all mice of this group had died before reaching this timepoint. **B)** Clinical severity scores were determined for each mouse at 2, 24 and 48 hpi. A score of 0 denotes no clinical symptoms, a score of 5 denotes death. **C)** The blood was collected for bacterial titer determination. For the group infected with high inoculum of WT cells, no data was gathered at 48 hpi, because all mice of this group had died before reaching this timepoint. **D)** The body weight of the infected mice was determined at 2, 24 and 48 hpi. Body weight was normalized to the weight at the beginning of the infection experiment. The legend is applicable to all panels. Statistical significance for the number of CFU recovered from the lungs and from the blood, and for the body weight was determined using an unpaired T-test. Statistical significance for the clinical severity scores was determined using a ranked Mann-Whitney U-test. Dashed lines denote the limits of detection at 2/24 hpi (short dashes), and 48 hpi (long dashes).

At the 24 hpi timepoint, the lower number of CFUs recovered from the lungs of mice infected with the Δ*ompR* mutant strain compared to the WT strain did not reflect the number of cells recovered from the blood, where similar numbers of CFUs were recovered for both strains (Figure 4C). In contrast, at the 48 hpi timepoint, more cells could be recovered from the blood of the mice inoculated with 10^5^ CFU of WT cells, than of the Δ*ompR* mutant strain. The deletion of *ompR* thus seemingly resulted in a decreased ability to establish lasting infections in the lungs and blood of the infected mice.

This decrease in virulence of the Δ*ompR* strain, as observed through the decreased number of CFUs recovered from the lungs and blood, and the decreased clinical severity and mortality, was also reflected by a decreased loss of body weight in the infected mice. Mice infected with the Δ*ompR* mutant strain lost less body weight over the course of the infection experiment (Figure 4D). These data suggest that the deletion of *ompR* results in a significant decrease in the virulence of *K. pneumoniae* ATCC 43816.

## Discussion

In the present study, we set out to investigate the role of the EnvZ/OmpR TCS in gene expression, infection, and virulence of *K. pneumoniae* ATCC 43816. We observed that deletion of *ompR* in *K. pneumoniae* ATCC 43816 does not lead to an appreciable loss of fitness *in vitro*. In addition, susceptibility to clinically relevant antibiotics also did not change. The transcriptional profiling showed changes in transcriptional activity of several genes including virulence-associated genes (in *K. pneumoniae*), and antibacterial response genes (in human lung epithelial cells). The Δ*ompR* mutant was however heavily attenuated in a murine infection model. These data suggest that OmpR is an important regulator of virulence, and that it plays a crucial role in the ability of *K. pneumoniae* to establish invasive disease.

On the pathogen side, the deletion of *ompR* led to changes in expression of orthologous genes previously described to be controlled by OmpR in other species (including *dtpA*, *ompK35, ompC* (commonly known as *ompS1*)*, mrkA*, and *mglB)*, but also of genes outside of this previously established orthologous regulon. The gene commonly annotated as *ompC* in *E. coli* and *Salmonella* species was not significantly differentially expressed. The limited overlap between the OmpR regulons of *K. pneumoniae* and other Enterobacteriaceae is not unsurprising, as the OmpR regulons between *E. coli* and *Salmonella enterica* subspecies Typhimurium serovar Typhimurium, a Gram-negative bacterium more closely related to *E. coli* than *K. pneumoniae*, also differ substantially (13, 17, 20–22, 26, 27). As we could not identify the preferential DNA sequence for the binding of OmpR by comparing the sequences preceding the genes that had been identified to be differentially expressed, future studies using ChIP-seq may elucidate the preferential DNA sequence bound by OmpR, and its exact regulon in *K. pneumoniae*.

Through our dual RNA-seq experiment, we observed downregulation of genes involved in virulence in the *ompR* deletion strain. Interestingly, we observe the downregulation of *rmpC* (annotated here as IT767_02620), which is involved in the regulation of capsule production in *K. pneumoniae*. This observation contrasts with earlier work, that shows that deletion of *ompR* does not change the transcriptional activity of the *rmpADC* operon (30). Importantly however, differences in experimental setup between our work (host-pathogen setup, cell culture medium) and the earlier performed work (Colombia blood agar plate), may explain these differences. We also identified that a homologue of ferritin-like protein FtnB (IT767_03325) was lower expressed, which has been described to be important in virulence in *Salmonella* (50). Lastly, the observed lower expression of *mrkA*, encoding a fimbrial type 3 subunit, potentially results in reduced presence of type 3 fimbriae. Together with the observed loss of hypermucoviscocity (30), these results could explain the attenuation of the mutant strain.

On the host side, no individual antibacterial response genes were significantly differentially expressed. However, GO term GSEA results do suggest that the host cells mount increased antibacterial responses typical for epithelial cells (increased antimicrobial peptide production, increased innate immune responses, and increased neuronal communication) towards the mutant strain than towards the WT strain (51). The increased expression of these groups of genes suggests enhanced clearance, and could explain the reduced virulence of the *ompR* knockout strain in the mouse infection model.

Even though we have identified the possible mechanisms through which a deletion of *ompR* may lead to reduced virulence *in vivo*, further experimentation may lead to a deeper biological understanding. The lack of single gene-level evidence may result from the fact that only two samples were available for the *ompR* deletion mutant, impacting statistical power. In addition, the overwhelming transcriptional response due to the presence of lipopolysaccharides (LPS), which are considered the prototypical pathogen-associated molecular pattern, might obscure any subtle host transcriptional effects from other missing epitopes because of the loss of OmpR. In any case, it seems that *ompR* is not crucial during the early stages of lung infection but is rather required for survival at later stages.

In our experiments, we confirm the previously reported reduced ability to establish an invasive infection of a OmpR-deficient ATCC 43816 strain. Fewer CFUs were recovered from the lungs and blood of mice infected with the Δ*ompR* strain, than from WT-infected mice at 48 hpi. In addition, these mice had a decreased mortality, lost less body weight, and had decreased clinical severity scores. Interestingly however, the number of WT and Δ*ompR* cells that was recovered from the blood at 24 hpi was not significantly different, in contrast to the number of CFUs recovered from the lungs at that timepoint. This suggests that loss of OmpR does not affect the ability of *K. pneumoniae* to disseminate from the lungs to the blood. However, OmpR does seem crucial for *K. pneumoniae* for survival in both blood and lungs.

Because of the central role of TCSs in bacterial adaptation to specific environmental cues, the development of inhibitory drugs against these systems has been explored before (9, 29). Our and work performed by others previously suggests that, because of the key role of EnvZ/OmpR in pathogenesis, newly developed anti-infective compounds targeting EnvZ/OmpR would be potent drugs to treat infections with a low potential for drug resistance development since bacterial growth is not perturbed *per se*. Although a decrease in susceptibility to tetracyclines was noted, tetracyclines are rarely used clinically to treat infections with *K. pneumoniae*. The deletion of *ompR* did not change the susceptibility to antibiotics typically used to treat infections with *K. pneumoniae* in clinical cases, including cephalosporins and carbapenems. The development of anti-virulence drugs targeting TCSs might thus be a fruitful approach to treat infections with multidrug-resistant bacteria.

## Author Contributions

A.B.J. conceived, designed, and performed experiments, analyzed data, and wrote the first draft of the manuscript. V.d.B. analyzed data, and edited the manuscript. R.A. conceived, designed, performed experiments, and analyzed data. V.T. conceived, designed, and performed experiments, analyzed data, and edited the manuscript. C.K. conceived and designed experiments, and supervised the study. M.P. conceived and designed experiments, supervised the study, and edited the manuscript. J.-W.V. conceived and designed experiments, and supervised the study.

All authors reviewed and approved the final version of the manuscript.

## Data availability

The raw Illumina sequencing reads of the transcriptomics data set are available in Gene Expression Omnibus (GEO) under accession number GSE123964 (https://www.ncbi.nlm.nih.gov/geo/query/acc.cgi?acc=GSE1239640).

## Acknowledgements

We thank V. Benes (GeneCore, European Molecular Biology Laboratory, Heidelberg) for his continuing support in library preparation and sequencing. Additionally, we would like to thank C.R. Aranzamendi Esteban and S. El Aidy for their access to human cell culture laboratory. We thank L. Ferrari and A. Felici at Aptuit (today Evotec) for the planning, conducting, and analysis of the mouse experiments.

