## Supplementary Information for "*Klebsiella pneumoniae* OmpR facilitates lung infection through transcriptional regulation of key virulence factors"

### Supplemental Figures

#### Supplemental Figure S1

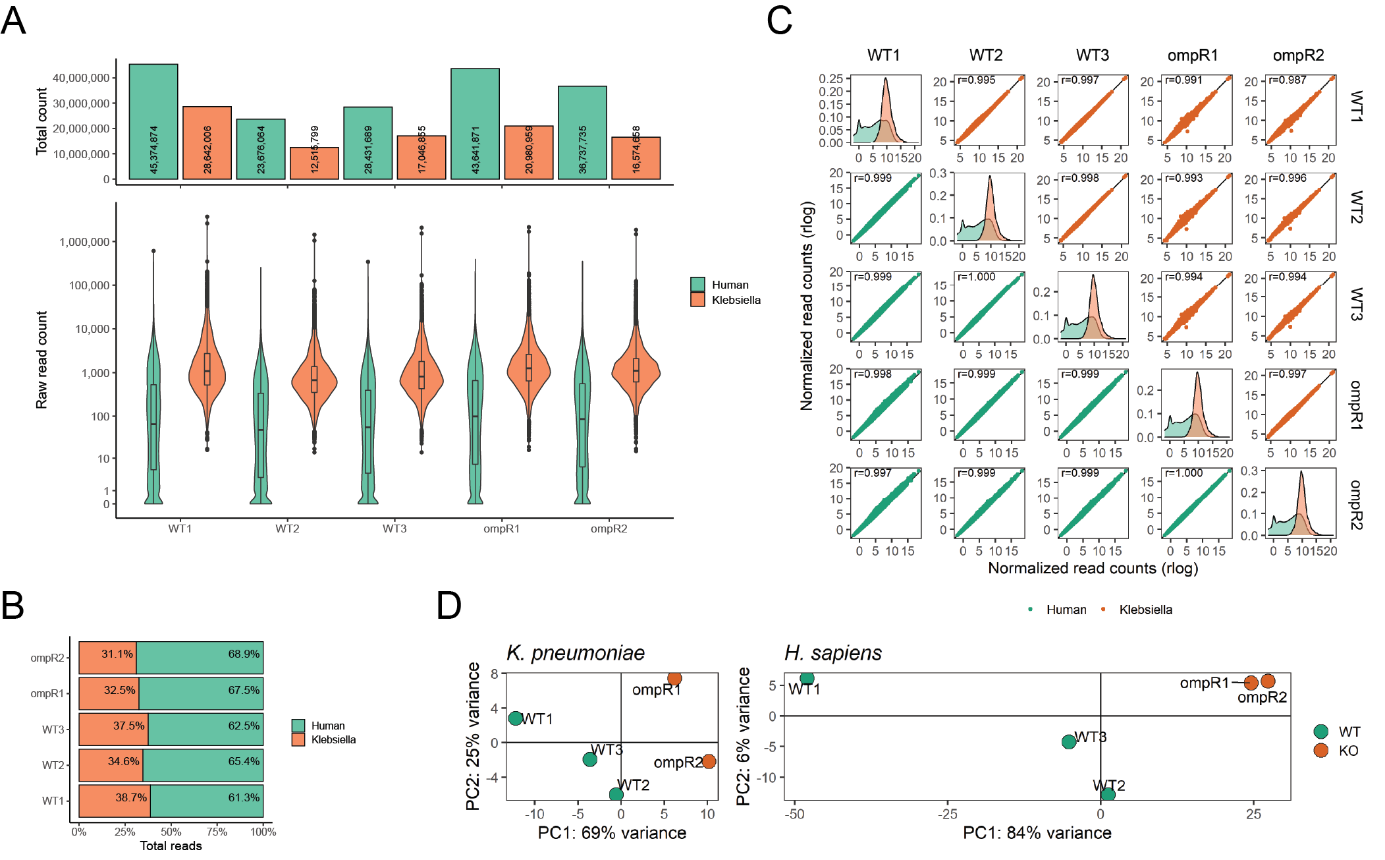

#### Supplemental Figure S2

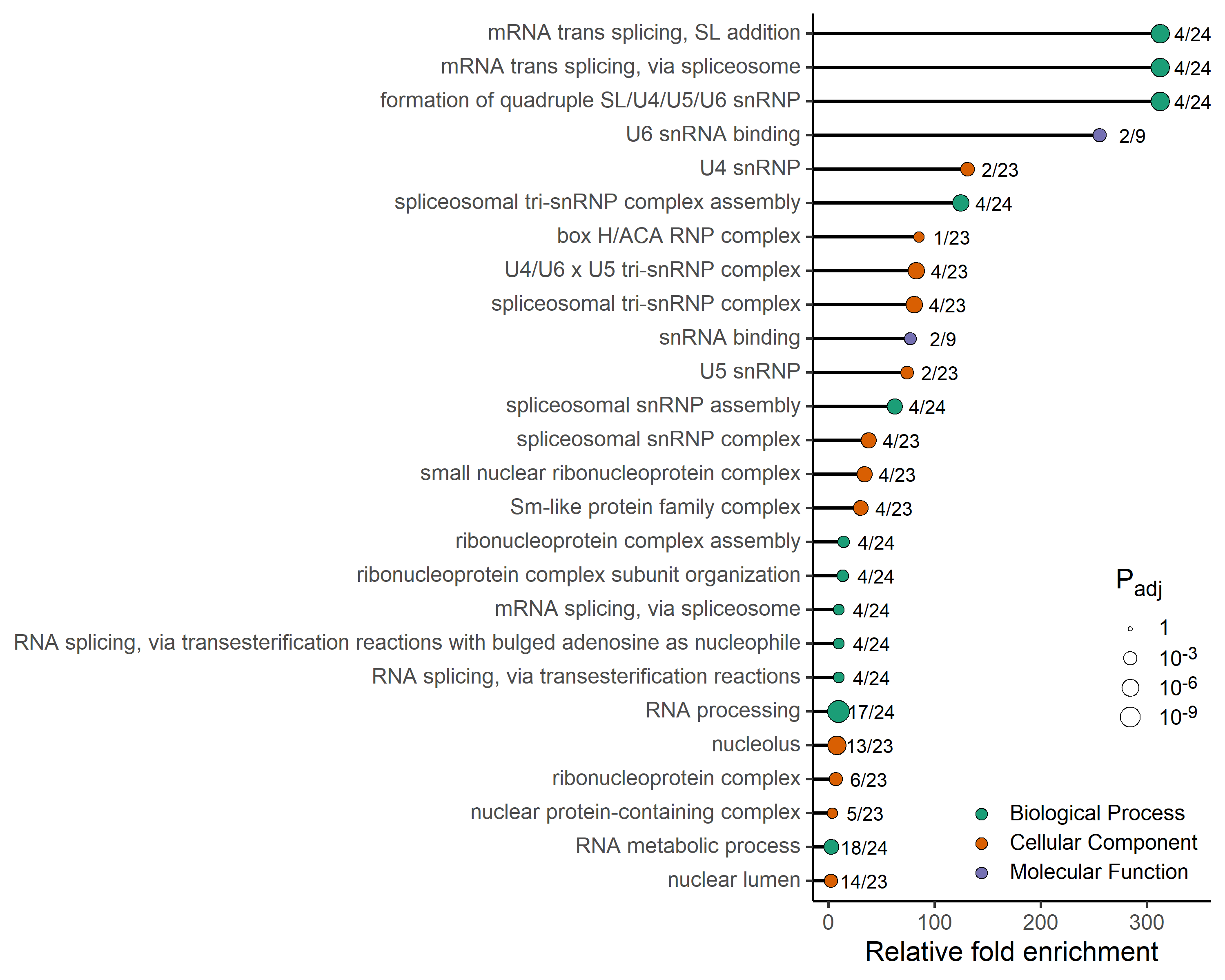

### Supplemental Figure Legends

#### Supplemental Figure S1

**Composition and mapping of dual RNA‑seq Illumina sequencing reads.** **A)** Distribution of the aligned and counted sequencing reads between the five libraries, split out per organism. The samples had an average of 54.7 million reads (range: 36.2 to 74.0 million reads) per library. **B)** The proportion of the sequencing reads of each sample mapped to the *K. pneumoniae* ATCC 43816 genome, and the human genome. **C)** Intra‑library distributions and inter‑library correlations of normalized counts. Read counts were normalized with a blind rlog transformation using DESeq2. **D, E)** Principal component analyses (PCA) plots of read counts for *K. pneumoniae* (panel D) and *Homo sapiens* (panel E).

#### Supplemental Figure S2

**GO term enrichment analysis of significantly downregulated genes in lung epithelial cells in response to Δ*ompR K. pneumoniae***. Relative fold enrichment is the gene ratio (faction of downregulated genes in GO gene set divided by downregulated genes in GO data base) divided by the background ratio (fraction of total genes in GO set divided by total genes in GO data base). Ratios displayed in plot are gene ratios. All significantly enriched GO terms are shown (P_adj_<0.05). Complete table of results can be found as Supplemental Table S5.

### Supplemental Tables

#### Supplemental Table S1

*See appendixes for full results of MIC determinations.*

#### Supplemental Table S2

*See appendixes for full DESeq2 results of Klebsiella pneumoniae samples.*

#### Supplemental Table S3

*See appendixes for full DESeq2 results of human lung epithelial cell samples.*

#### Supplemental Table S4

*See appendixes for full results of GO term enrichment analysis of significantly differentially expressed genes in the human lung epithelial cells.*

#### Supplemental Table S5

*See appendixes for full results of GO term enrichment analysis using all available log2FC values of genes in the human lung epithelial cells.*

### Supplemental Table Legends

#### Supplemental Table S1

**MIC values for ATCC 43816 WT and ATCC 43816 Δ*ompR***. Minimal inhibitory concentrations of several types of antibiotics were determined for both the ATCC 43816 WT and the ATCC 43816 Δ*ompR* strain, using microbroth dilution methods (except for the MIC of ertapenem for the WT strain, which was determined on a Vitek 2). All MIC values are given in µg/ml.

#### Supplemental Table S2

**Full results of DESeq2 analysis of the genes in the *K. pneumoniae* genome.** Complete standard DESeq2 differential expression analysis output plus gene annotation details. Hypothesis tests were performed against an alpha of 0.05 and log_2_FC threshold of log_2_(1.5): 1.5× mRNA depletion/enrichment in the Δ*ompR* bacteria compared to WT. The log_2_FC values were shrunk with the apeglm method.

#### Supplemental Table S3

**Full results of DESeq2 analysis of the genes in the human genome.** Complete DESeq2 differential expression analysis output, including gene annotation details. Hypothesis testing was performed with default parametrization: against an alpha of 0.01 and log_2_FC threshold of log_2_(1). The log_2_FC values were shrunk with the apeglm method.

#### Supplemental Table S4

**Full results of the GO term enrichment analysis for differentially expressed genes.** Standard clusterProfiler output for all three GO data bases, based on shrunk log_2_FC values of the genes that were significantly differentially expressed, between the human lung epithelial cells challenged with Δ*ompR* *K. pneumoniae* and those challenged with WT *K. pneumoniae*. The log_2_FC values were shrunk with the apeglm method as implemented in DESeq2.

#### Supplemental Table S5

**Full results of the GO term gene set enrichment analysis.** Standard clusterProfiler output for all three GO data bases, based on shrunk log_2_FC values of human lung epithelial cells challenged with Δ*ompR* *K. pneumoniae* and those challenged with WT *K. pneumoniae*. The log_2_FC values were shrunk with the apeglm method as implemented in DESeq2.
